# Mitigation of membrane morphology defects explain stability and orientational specificity of CLC dimers

**DOI:** 10.1101/2023.03.16.533024

**Authors:** Tugba N. Ozturk, Nathan Bernhardt, Noah Schwartz, Rahul Chadda, Janice L. Robertson, José D. Faraldo-Gómez

## Abstract

Most membrane proteins are oligomers, but the physical forces explaining the stable association of these complexes inside the lipid bilayer are not well understood. The homodimeric antiporter CLC-ec1 highlights the puzzling nature of this reaction. This complex is thermodynamically stable even though it associates via a large hydrophobic protein-protein interface that appears well adapted to interact with the membrane interior. In a previous study, however, we discovered that when CLC-ec1 is dissociated, this interface introduces a morphological defect in the surrounding membrane, leading us to hypothesize association is driven by the elimination of this defect upon dimerization. This study tests this hypothetical mechanism directly and shows it is supported by molecular and physical models. First, using coarse-grained umbrella-sampling molecular simulations, we calculated the membrane contribution to the potential-of-mean-force for dimerization in a POPC bilayer. This shows the stable association of CLC subunits prior to formation of direct protein-protein contacts, but only via the native interface that presents the membrane defect, and not others. Single-molecule photobleaching experiments show that addition of short-chain DLPC lipids, known to alleviate the membrane defect, also shifts the association equilibrium from dimers to monomers. We explain this destabilizing effect through additional umbrella-sampling and alchemical free-energy simulations, which show DLPC enrichment of the defect diminishes the membrane contribution to the association free energy, as it improves the lipid-solvation energetics of the monomer but not the dimer. In summary, this study establishes a physical model that explains the stability and orientational specificity of CLC dimers in terms of membrane-mediated forces, rather than protein-protein interactions. We posit that cells might ubiquitously leverage morphological defects in the bilayer to drive organization of membrane proteins into functional complexes, and that cellular regulation of lipid composition can modulate this organizing effect.

## Introduction

Membrane proteins spontaneously self-assemble into folded subunits, oligomeric assemblies and higher-order complexes in membranes. This is apparent by the vast number of examples of membrane protein structures, oftentimes oligomers, present in the Protein Data Bank (Aleksandrova, Sarti, and Forrest 2020). Yet, the physical reasons for how stability is achieved in membranes, and the factors that modulate the stability, remain unclear due to the limited amount of experimental data on free energies of assembly in membranes. From the data available to us, strong affinity has been observed in complexes such as glycophorin-A dimers largely attributed to backbone hydrogen bonding and van der Waals interactions (Smith et al. 2002; Mueller, Subramaniam, and Senes 2014). However, other membrane protein complexes are predominantly hydrophobic, lacking contenders for hydrogen bonding. For surfaces that bind via side-chain contacts, the role of van der Waals interactions remains in question, as similar van der Waals interactions can be achieved with solvating lipids. With this, the question remains as to whether there is a different, generalizable physical driving force that arises from the membrane to non-specifically promote the folding and assembly of membrane proteins. Such a physical force would be similar in essence to the hydrophobic effect being a common driver for soluble protein folding and oligomerization that is also generalizable in that it does not rely on specific protein sequence. In the membrane, it is conceivable that such a driving force could enable protein organization and ordering, even at a distance, as observed in some systems (Anselmi, Davies, and Faraldo-Gómez 2018). Finally, a common membrane-dependent driving force provides a rationale for the vast complexity of membrane environments that are observed across biology, which could play a physical role in the membrane protein reactions within. To understand the driving force for protein association in membranes, and whether it is membrane-dependent, we must have a way of analyzing these reactions in thermodynamically meaningful ways by partnering experiments with molecular and physical models.

Recently, our group developed a model system where reversible, equilibrium dimerization within the membrane can be experimentally quantified to measure the thermodynamically meaningful standard state free energy of dimerization. In this approach, single molecule photobleaching analysis of equilibrium subunit capture is carried out to quantify redistribution of protein populations within the membrane while controlling for thermodynamic variables. These studies have been applied to several systems, including the homodimeric chloride/proton antiporter CLC-ec1 (**Fig. 1A**) from *E. coli* (Chadda et al. 2016)and the dual-topology homodimeric fluoride ion channel Fluc (Ernst et al. 2023). But CLC-ec1 remains a particularly interesting system in that it forms dimers via a membrane embedded, hydrophobic interface, achieving strong stability in biologically relevant membranes, with *ΔG°*_*2:1POPE/POPG*_ = -10.9 kcal/mole (Chadda et al. 2018) and *ΔG°*_*EPL*_ = -12.7 kcal/mole (Chadda et al. 2023), relative to the 1 subunit/lipid standard state. This is a biologically stable form, as 2 copies in the *E. coli* cell would lead to 97% in the dimeric form at any given time, suggesting that the physical mechanism defining the stability is achievable even though it occurs at a hydrophobic interface. Our recent study on CLC showed that the membrane structure in the monomeric state is significantly perturbed, forming a non-bilayer defect that is thinned and twisted, leading to significant lipid tilting and a decrease of density that reduces packing and increases water penetration. This is in line with more recent measurements that indicate that the thermodynamic basis of dimerization is accompanied by a large negative change of heat capacity, indicative of ordering of lipids and water in the monomeric state (Chadda et al. 2023). Furthermore, we find that addition of a better solvent for this defect, specifically short-chain di-lauryl C14 lipids, both improve lipid packing in the solvation structure, and is experimentally capable of shifting the equilibrium towards monomers entirely, while preserving transport function. Finally, the linkage of the change in equilibrium with short-chain lipid activity demonstrates a mechanism of preferential solvation, which we also observe in our co-mixture simulations with short chain DL lipids preferentially enriched in the membrane defect. Altogether, these results strongly suggest that the innate driving force for CLC dimerization comes from the gain in free energy from eliminating the membrane defect in the dimer state. In this case, dimerization can be driven by the membrane energetics alone where the protein structure plays the role of defining the membrane defect.

**Figure 1.**
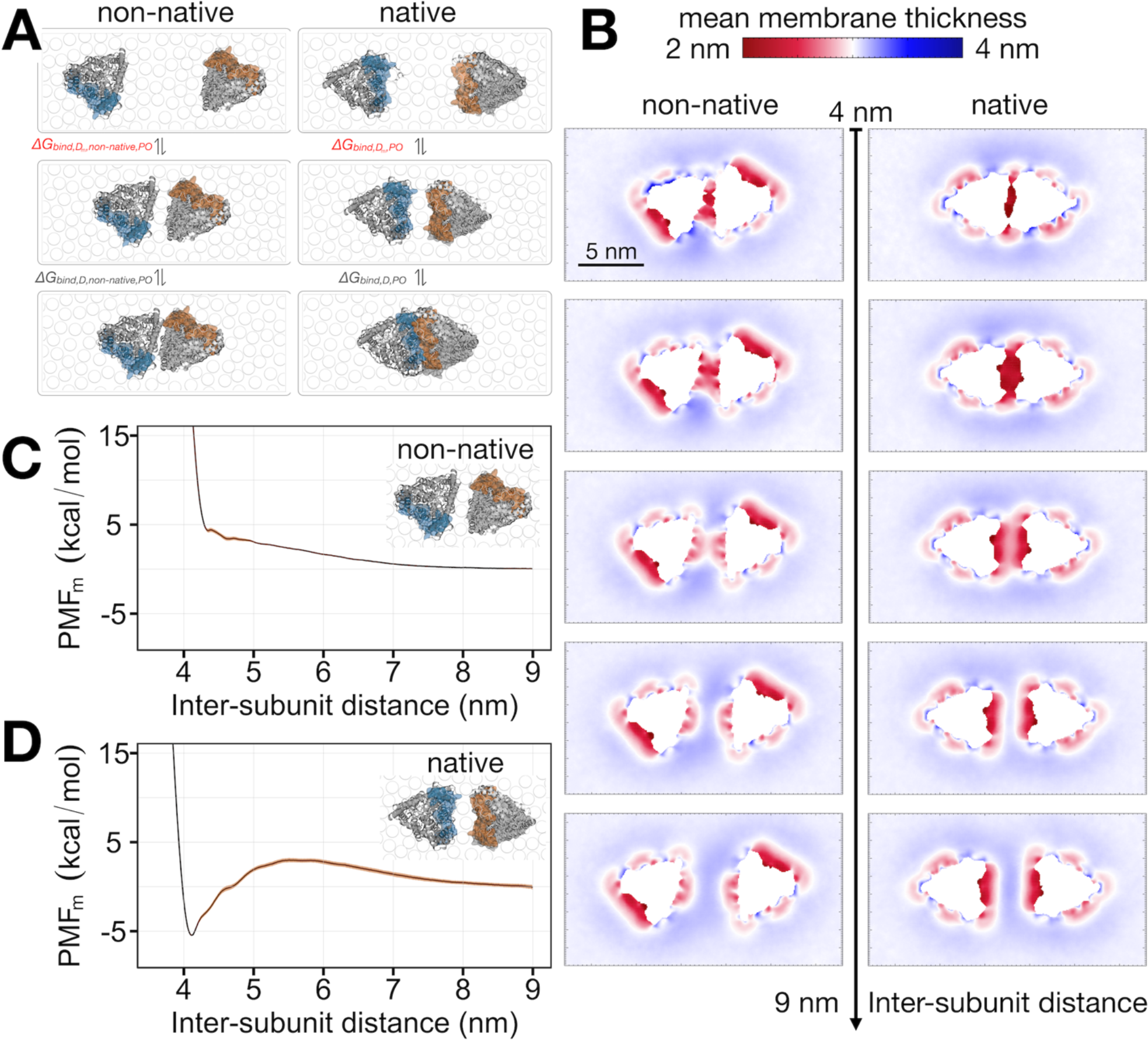
The membrane-dependent driving force for CLC-ec1 dimerization. **(A)** The dimerization reaction of the non-native CLC dimer complex generated in HDOCK (*left*) and native dimer complex (*right*) based on pdb 1OTS(Dutzler, Campbell, and MacKinnon 2003). The free energy of dimerization is separated into two steps, *ΔG*_*dimerization*_ = *ΔG*_*bind,Dm*._ *+ ΔG*_*bind,D*_, where D_m_ represents the dimer state where the two subunits come close but do not form direct contacts. We define the reaction between the dissociated monomers and the non-contact dimer as providing the membrane dependent contribution to the free energy, and the potential of mean force from membrane dependent driving forces **(B)** Average membrane thickness maps calculated for non-native and native subunit configurations between the dimer state and 5.5 nm separation representing dissociated monomers, over 10 μs simulation time. **(C)** Potential of mean force profile for non-native CLC-ec1 dimerization in membranes, isolating the membrane contribution (PMF_m_). **(E)** PMF_m_ for dimerization of CLC-ec1 at the native dimerization interface. Shaded orange bands represent the standard error values around the mean PMF values estimated with block averaging from four separate parts of these trajectories.

To test this theory, we examine whether this hypothesis is supported by molecular and physical models by carrying out free energy calculations of the dimerization reaction considering the membrane-only contribution. Since the major limitation in modeling membrane reactions is sampling of the lipid components, we choose the coarse-grained Martini force-field combined with umbrella sampling across the inter-subunit distance coordinate. By excluding configurations containing protein contacts, we calculate the membrane-only contribution to the free energy profile and find that dimerization is energetically favored only at the native dimerization interface where the membrane defect occurs. Next, we calculate the change in free energy upon adding short-chain lipids, by calculating the potential of mean force in the mixed lipid condition, as well as the change in the solvation free energy association with adding short-chain lipids using an alchemical free energy perturbation approach. Both computational approaches demonstrate that the addition of DL lipids destabilizes dimerization, in agreement with our experimental studies. Therefore, our results demonstrate that the CLC dimerization is driven by the change in the solvation free energy in the dimeric and monomeric states, and the magnitude of this driving force can be tuned by altering the chemical composition of the membrane.

## Results

### The membrane contribution to the potential-of-mean-force for CLC dimerization

In our previous study we postulated that the association of CLC-ec1 subunits is driven by a gain in free energy achieved by relieving membrane defects upon burying the perturbative and highly complementary dimerization interface (Chadda et al. 2021). Here, we examine whether this hypothesis is physically supported by carrying out a free energy calculation for the dimerization reaction, designed to isolate the contribution that comes from the membrane alone. To do so, we first used the approach of umbrella sampling. This method allows for focused sampling across a specific reaction coordinate, which we define as the inter-subunit distance of the two protomers. Harmonic potentials are applied in 0.5 Å separation increments, starting with the minimum inter-subunit distance defined by dimer complex and up to a final separation of 5.5 nm for a total of 111 simulation windows. To focus our sampling further, we apply a series of limited orientational restraints to ensure that the binding interfaces of interest are always oriented *en face*, while still allowing subunits to wobble relative to each other. Finally, since we anticipate large scale changes in membrane structure and a need to sufficiently sample lipid exchange in each configuration, we choose to use the coarse-grained Martini force-field that allows us to maximize the sampling of these factors. While this force-field represents a reduced physical model, we can examine robust contributions from the membrane and their role in defining the free energy changes associated with this reaction.

Before carrying out the potential of mean force (PMF) calculation for association at the native CLC dimerization interface, we sought out a negative control that would provide a reference for our calculation approach. CLC has only been observed to dimerize at the native interface and is not expected to interact stably at its other interfaces. Based on this, we built a non-native CLC-ec1 dimer model using the HDOCK server (Yan et al. 2020). In this model, instead of binding occurring at helices H, I, P and Q, the subunits are rotated relative to each other such that dimerization involves helices B, C and F (**Fig. 1A, Supp. Fig. 1)**. Notably, there is no significant membrane thinning at the non-native interface as it is typically exposed to the membrane and hydrophobically matched to the surrounding membrane thickness (**Fig. 1B**). We carried out umbrella sampling of the dimerization reaction at this non-native interface, and calculated the membrane-only contribution to the PMF using the force-correction analysis method (FCAM) (Marinelli and Faraldo-Gómez 2021). FCAM generates the PMF profile after distributing the trajectory frames into individual bins and estimating the mean force of the system in response to the applied bias potential at each bin. To calculate the membrane contribution, a *post hoc* filtering can be applied to the PMF calculation to exclude trajectory frames that contain protein-protein contacts. In this case, the PMF was calculated with frames containing less than 15% of the inter-subunit contacts that are observed in the original non-native homodimer model. This yields an estimate of the free energy contribution that does not involve significant protein-protein interactions, which we define as the membrane contribution, denoted as PMF_m_ (**Fig. 1C**). With this approach, we find that the PMF_m_ for the non-native complex is unfavorable, showing a repulsive potential as the two protein subunits come closer together but before they make direct contact. Therefore, we find that there is no membrane dependent driving force for CLC dimerization at this non-native interface.

Next, we calculated whether there was a membrane dependent driving force for dimerization at the native and conserved CLC binding interface. We carried out the exact same strategy of umbrella sampling across the inter-subunit distance reaction coordinate as we did for the non-native control, but now with the native dimerization interfaces oriented towards each other such that the membrane defects were aligned. Examining the mean membrane thickness maps as a function of the inter-subunit separation shows how the two defects coalesce into one as the two subunits come closer together (**Fig. 1B**). In contrast to the findings for the non-native complex, the PMF_m_ for binding at the native interface is clearly favorable, with stabilization occurring once the two membrane defects begin to merge, without direct protein-protein contacts (**Fig. 1D**). Furthermore, the stabilization increases as the defect area gets smaller, indicating that it is the removal of the membrane defect that is the stabilizing factor, demonstrating a membrane-dependent driving force for CLC dimerization within the lipid bilayer.

### CLC-ec1 dimerization is experimentally destabilized in short-chain PC lipids

Previously, we showed that the addition of short-chain C14:0 di-lauryl (DL) lipids to a 2:1 POPE/POPG, C16:0/18:1 membrane shifts the CLC-ec1 dimerization equilibrium to the monomeric form in a concentration dependent manner (Chadda et al. 2021). While we can only measure relative changes in the free energy, the most likely cause for this is a change in the solvation free energy of the monomeric state, as the shorter DL lipids pack better in the thinned membrane defect that forms around the exposed dimerization interface (Chadda et al. 2021). Supporting this theory, we computationally observed the preferential enrichment of DL lipids around the exposed dimerization interface in the monomeric state, indicating that DL partitioning into the membrane defect is energetically favorable. Altogether, these experimental and computational studies provide further support for a membrane-dependent mechanism of CLC dimerization, as it indicates that the driving force can be tuned by reducing the energetic penalty with changing the chemical composition of the solvent.

To test this hypothesis further, we carried out a new set of experiments to examine the extent of this acyl chain length dependency on CLC-ec1 dimerization stability. If the solvation free energy of the thinned membrane defect depends on improved acyl chain packing, then we expect that this effect will persist even in membranes containing different headgroup chemistry. To examine this, we measured the CLC monomer-dimer distributions in POPC and with the addition of DLPC using the single-molecule photobleaching subunit-capture approach (Chadda et al. 2016; Chadda and Robertson 2016). While the photobleaching probability distributions obtained in 2:1 POPE/POPG and POPC membranes were similar and indicative of a dimeric population, addition of 20% DLPC shifted the population significantly to a monomeric distribution, indicated by the increased observation of single photobleaching steps (**Fig. 2A**). Since an increase in monomers could also indicate non-reactive states such as misfolded subunits, we tested the CLC proteoliposomes for chloride transport function in the different membrane conditions. In fact, transport function increased in the 20% DLPC, 80% POPC membranes compared to the 100% POPC condition, where we observed the protein to be dimeric (**Fig. 2B-D**). Therefore, we conclude that the increased probability of single photobleaching steps corresponds to the presence of functional CLC monomers in the membrane. Overall, these results show that the equilibrium of CLC dimerization is dependent on the lipid composition, but specific to the acyl-chain length and not headgroup chemistry, in line with lipid packing defining the energetic cost of solvating the monomeric state.

**Figure 2.**
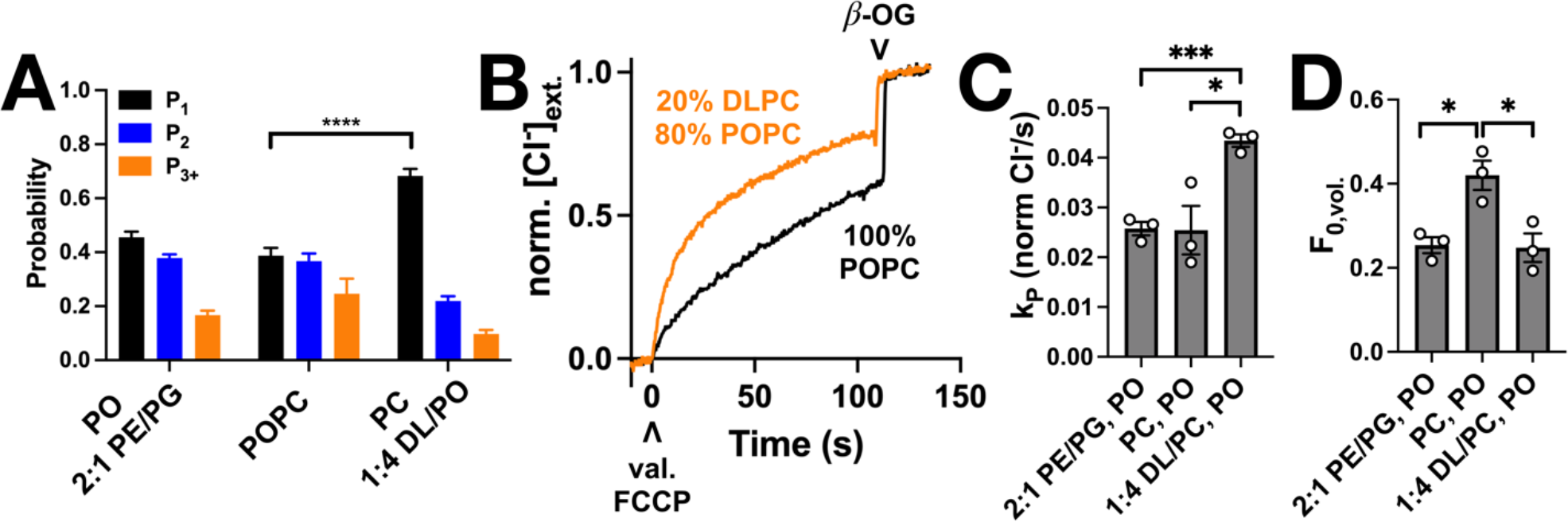
CLC dimerization is destabilized in 20% DLPC, 80% POPC membranes while transporter activity is enhanced. **(A)** Single-molecule photobleaching probability distributions of CLC-ec1-Cy5 0.1 µg/mg, *χ*_*reconst*._ = 10^−6^ subunits/lipid in 2:1 POPE/POPG, POPC and 1:4 DLPC/POPC membranes. Data represented as mean ± sem for n = 3-9 independent samples imaged 2-5 days after freeze/thaw fusion. A *χ*^2^ test was carried out on the different distributions: ****, *p* < 0.0001; not significant otherwise, *p* > 0.05. **(B)** Raw chloride efflux trace from a 1 μg/mg, *χ*_*reconst*._ = 10^−5^ subunits/lipid reconstituted in POPC (black) or 1:4 DLPC/POPC (orange). The upward caret (^) indicates initiation of transport with valinomycin and FCCP, whereas the downward caret (v) indicates addition of detergent β-OG releasing the remaining trapped chloride within the proteoliposomes. **(D)** The rate of chloride transport, *k*_*P*_ (normalized Cl^-^/sec) and **(C)** fraction of inactive vesicles, *F*_*0,vol*_. Data represent mean ± SEM, n = 3-5. Unpaired t tests were carried out between DL vs. PO conditions: *, *p* < 0.05; ***, *p* < 0.001.

### Addition of short-chain lipids destabilizes dimerization by changing membrane energetics

Next, we examined whether the lipid dependency of the CLC dimerization free energy is supported by our physical model. Here, we carried out the same umbrella sampling approach to calculate the membrane dependent PMF_M_, but now in 30% DLPC, 70% POPC lipids (**Fig. 3A**). Since this is now a more complex binary lipid system, we increased sampling times to 40 μs per window to reach convergence. Despite the overall 1.6 Å thinning of the bulk membrane, we found similar thinned membrane defects around the dimerization interface (**Fig. 3B**) as we observed in our previous simulations of DL/PO systems (Chadda et al. 2021). In addition, we also find that DL lipids become enriched in the membrane defect (**Fig. 3C**). Finally, we observe the water penetration into the hydrophobic core of the POPC membranes, is reduced in the mixed DL/PO bilayers (**Fig. 3D**). In line with these molecular observations, we find that the PMF_m_ for CLC dimerization in 30% DLPC, 70% POPC is significantly less stable than in 100% POPC membranes (**Fig 3E**), in line with a stabilization of the solvation structure in the monomeric state. A rough estimate of the magnitude of the destabilization is obtained by taking the difference in the energetic wells corresponding to when the two defects coalesce, yielding *ΔΔG*_*PMF*,0→30%DL_ = +1.9 ± 0.5 kcal/mole at the well minima. This value is in qualitative agreement with the experimentally measured changes in dimerization free energy for CLC-ec1 in 2:1 DLPE/DLPG and 2:1 POPE/POPG membranes.

**Figure 3.**
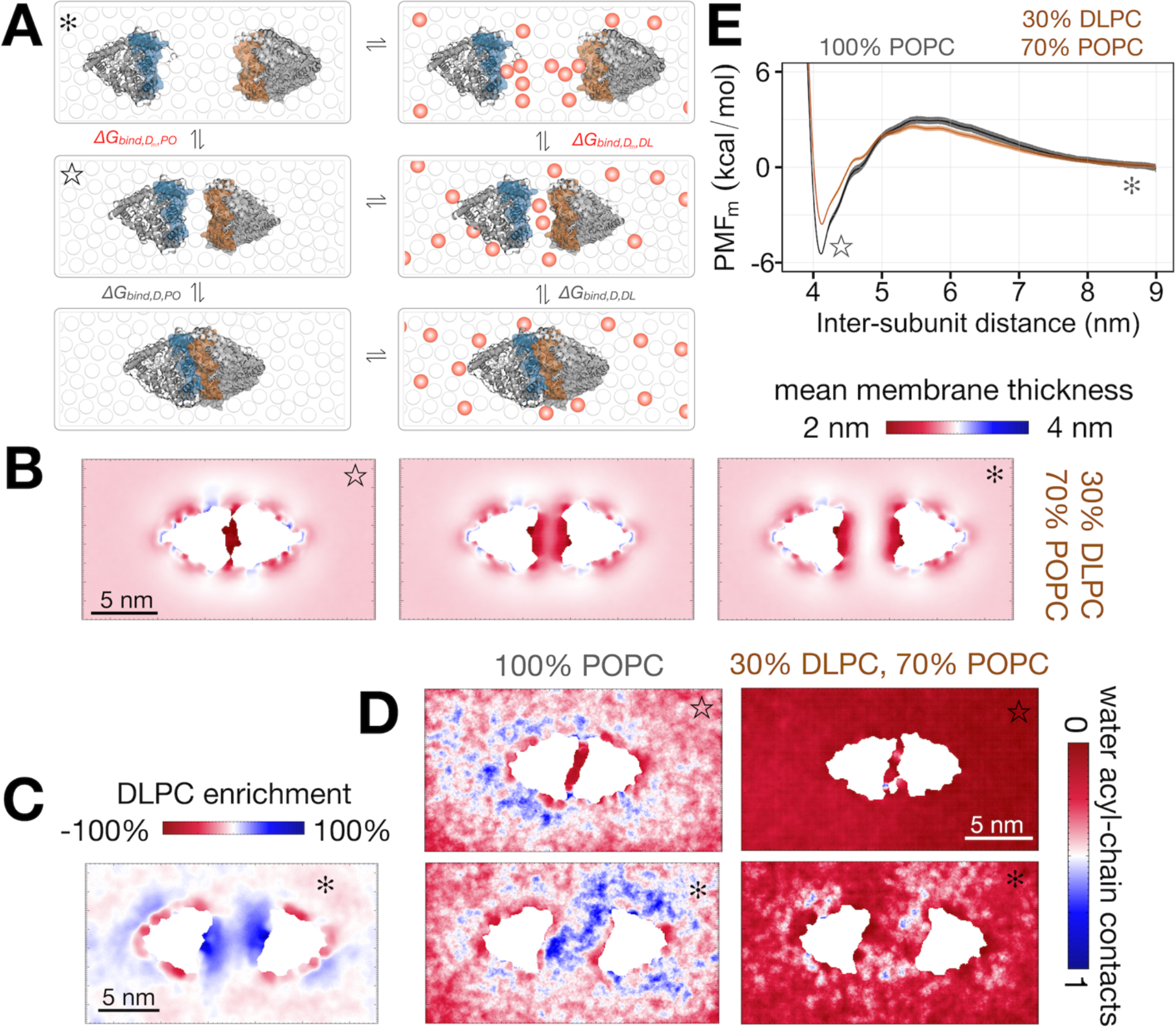
The PMF_m_ for CLC dimerization is destabilized by short-chain lipids. **(A)** The change in free energy for CLC dimerization in the native complex as a function of lipid composition is determined by calculating one PMF_m_, in a reference lipid, e.g. POPC, and one in the test lipid, e.g. 30% DLPC, 70% POPC. The difference in the free energy due to the change in lipid composition is *ΔΔG*_*PMFm*,0→30%DL_ = *ΔG*_*PMF,Dm,DL*_-*ΔG*_*PMF,Dm,PO*_, where each term is calculated from the respective PMF_m_. **(B)** Two-dimensional maps of mean membrane thickness computed over the 40 µs MD simulations. **(C)** Percent enrichment of DLPC lipids in the lower leaflet. **(D)** 2D maps of the number of contacts between water and lipid tails in the lower leaflet in 100% POPC (10 *μ*s) and simulations from the native model in a 30% DLPC, 70% POPC (40 *μ*s). **(E)** PMF_m_ for 30% DLPC shows that CLC dimerization is energetically destabilized compared to the POPC condition, with *ΔΔG*_*PMFm*,0→30%DL_ = 1.9 ± 0.5 kcal/mole at the well minima. Shaded regions represent the standard error calculated using potential of mean force profiles generated using four even blocks of the 10 *μ*s-long trajectories of umbrella sampling windows for 100% POPC and 40 *μ*s-long trajectories in 30% DLPC, 70% POPC.

This change in the membrane dependent component of the dimerization free energy, *ΔΔG*_*PMF*, 0→30%DL_, is equivalent to the change in the solvation free energy associated with change in lipid composition, *ΔΔG*_*solvation*,0→30%DL_, as they represent alternate legs of the same thermodynamic cycle (**Supp. Fig. 2)**. To calculate the change in the solvation free energy directly, we employ the strategy of alchemical free energy perturbation (FEP) method. In this approach, we consider the free energy difference of the thermodynamic cycle of the monomer and dimer states in two different lipid compositions (**Fig. 4A**). The free energy difference between these two legs of the cycle can be calculated by alchemically converting a certain number of POPC lipids to shorter DLPC lipids by free energy perturbation (**Fig. 4B**). Since the ultimate transition from 0→30% DLPC involves many molecules, we split the computation into multiple segments such that DLPC lipids are gradually added to the system (**Supp. Table 4**), which also allows us to compare with the experimental DL titrations. Four transformation steps are carried out, starting with 0→1%, 1→10%, 10→20% and finally 20→30% DLPC. Of course, the FEP methodology is built on the assumption that the phase space dimensions are unaffected by a given transformation, i.e., the number of atoms does not change. In cases where one or more atoms are deleted, this limitation may be circumvented by introducing dummy atoms whose through space interactions are muted. For our simulations, we created a modified DLPC* that is identical to the original DLPC but with a dummy atom attached to the end of each alkyl chain. MD simulations containing DLPC* are indistinguishable to those containing DLPC, as indicated by established metrics like membrane thickness and leaflet interdigitation (**Supp. Fig. 5**). Importantly, the DLPC* molecules preserve the spatial distribution of the lipids as can be seen by examination of the DLPC enrichment factor (**Supp. Fig. 7**).

**Figure 4.**
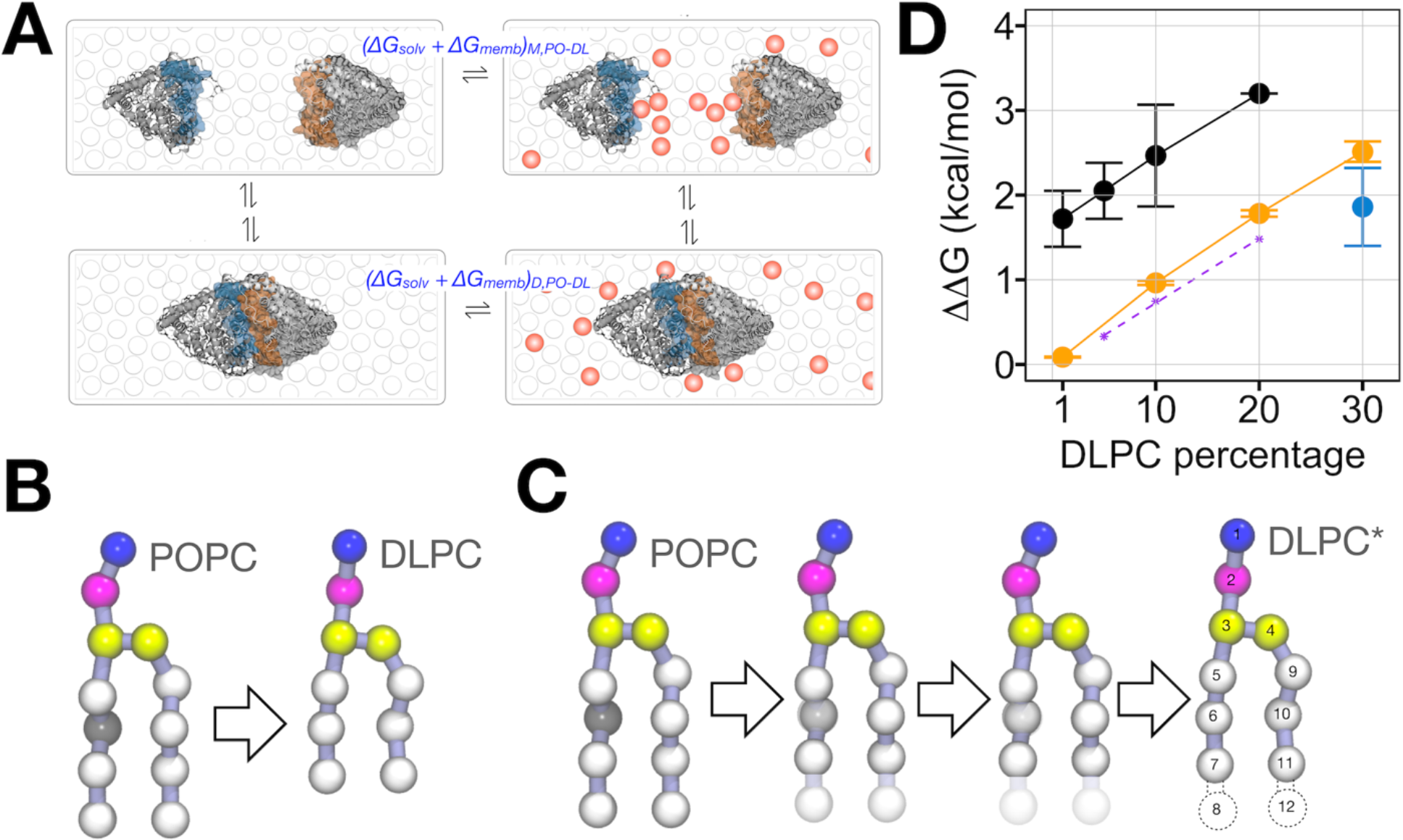
Free energy perturbation calculations for the change in CLC solvation free energy association with addition of DL to PO membranes. **(A)** The change in the solvation free energy due to change in the lipid composition is obtained from the vertical legs of the thermodynamic cycle (**Supp. Fig. 2**). **(B)** This can be obtained by carrying out an alchemical free energy perturbation calculation where a certain number of POPC lipids are converted to POPC in the following compositional increments: 0→1%, 1→10%, 10→20% and finally 20→30% DLPC. **(C)** To maintain the number of atoms in the system, the actual conversion involves transformation of POPC to DLPC*, representing a DLPC molecule with a dummy atom attached to each acyl chain that is otherwise transparent in all physical properties. Simulations of membranes containing DLPC* are identical to DLPC systems (**Supp. Fig. 5)** and are suitable for use with our FEP simulations. These simulations were constructed to minimize the relative entropy between neighboring replicas **(Supp. Fig. 6)** and consequently the error in each free energy difference, which is estimated from discrepancies between the forward and reverse pathways **(Supp. Fig. 7). (D)** *ΔΔG*_0→X%DL_ relevant to the CLC dimerization free energy due to the change in solvent composition. Experimental *ΔΔG*_*expt*.,0→X%DL_ data (black) (Chadda et al. 2021), *ΔΔG*_*PMF*,0→30%DL_ at the well minima (blue) and *ΔΔG*_*FEP*,0→30%DL_ (orange). *ΔΔG*_*expt*.,1→X%DL_ is designated by the dashed purple line.

Given the many similarities between DLPC* and DLPC, we transformed POPC molecules into DLPC* in our FEP simulations. These simulations were carried out using Hamiltonian replica exchange where the scaling parameter, lambda, was varied for each replica thus improving sampling. These simulations employed a set of lambda values derived by restricting the relative entropy between neighboring replicas to a value of 1 kT or less (**Supp. Fig. 6**). This requirement ensures significant overlap between the probability distributions of neighboring replicas and resulted in 38 replicas for the 0→1% transformation and 93 for the remaining (**Supp. Table 5**). To enable error estimates, we performed two independent simulations for each transformation capturing the forward and backward paths. That is, we performed a simulation where lambda was varied from 0 to 1 when moving across the replicas and another where this distribution was reversed. The estimated free energy cost for each transformation as a function of the simulation time was obtained using the Bennett Acceptance Ratio and are well converged within the course of each simulation (**Supp. Fig. 7**). Importantly, our simulations were long enough to relax the spatial distribution of the lipids as is seen by examination of the DLPC enrichment factor. We note that these distributions were slow to converge requiring 6 *μ*s for the 1→10%, 10→20%, and 20→30% transformations and 40 *μ*s for the 0→1% condition. In each case, *ΔΔG* is a small fraction of either *ΔG*, limiting the acceptable error for each measurement.

Upon summing the contribution from each step, we acquire *ΔΔG* for the complete path from 0 → 30% (**Fig. 4C**). With this, we find *ΔΔG*_*FEP*, 0→30%DL_ = 2.5 ±0.1 kcal/mole for the full transformation. This value is comparable to our expected value extrapolated from experimental measurements *ΔΔG*_*expt*.,0→30%DL_ ∼ 4.0 kcal/mole (Chadda et al. 2021) and the difference in free energy derived from the umbrella sampling calculations, *ΔΔG*_*PMF*,0→30%DL_ = 1.9 ± 0.5 kcal/mole. In addition, we also find that the slope of *ΔΔG* vs. % DL from the FEP calculations shows the same dependency as the experimental data. In fact, the only significant difference between the experiments and FEP calculations, is that we do not observe the abrupt shift in stability over the 0→1% change in DL, as observed in our experiments. Given the number of replicas used and the length of each simulation, we find it unlikely that this discrepancy stems from sampling deficiencies but may instead reflect a limitation of the system size and number of DL molecules in our system, or the coarse-grained Martini force-field.

## Discussion

This study presents physics-based evidence that CLC dimerization is driven by a membrane dependent change in the solvation free energy that is tunable by changes in lipid composition. The mechanism of CLC dimerization is in major part defined by how the dissociated monomer perturbs the surrounding bilayer structure. Since the dimerization interface is highly curved, and presents significant hydrophobic mismatch, the membrane must adjust to accommodate to the protein structure, leading to significant twisting, tilting of lipids and disruption of neighboring interactions that allow water to penetration into the hydrocarbon core. This perturbation comes with an energetic penalty, which is mitigated as the monomers associate, even before the two protein surfaces touch, as lipids in the defect return to the bulk lipid bilayer and reduce the free energy of the system. This implies that affinity by this mechanism, will be defined by the work required for the lipid bilayer to deform to optimally solvate the exposed dimerization interface. It will also depend on how effective dimerization hides this surface from the surrounding lipids, which will be a function of the protein surface complementarity. Our calculation of the membrane-only potential of mean force reveals that there is a favorable attraction that arises even when the two proteins do not touch, because the lipids are free from the obligation of solvating this interface. Note, our calculations showing the lack of stabilization at the non-native CLC surface indicates that this effect is specific, only occurring at interfaces that present significant perturbations to the membrane structure. This presents a general model for specific and stable membrane protein association in membranes by burial of membrane defects. This is a mechanism where exposed protein surfaces shape and deform the surrounding membrane away from the lipid bilayer structure, creating an intrinsic driving force to bury this specific interface in a manner that returns the most lipids back to the bulk membrane.

This model of membrane protein association by burial of defects also presents a mechanism for modulating protein association equilibria by changes in lipid composition. In this study, we demonstrate two independent methods of calculating the lipid dependency on protein association stability and find that the calculated *ΔΔG*_*PMF*,0→30%DL_ = 1.9 ± 0.5 kcal/mole and *ΔΔG*_*FEP*,0→30%DL_ = 2.5 ± 0.1 kcal/mole with our extrapolation of the experimental results, placing *ΔΔG*_*expt*.,0→30%DL_ around 4.0 kcal/mole. This is a major finding that offers insight into one reason for why biology may explore diversity in lipid chemistry, which may be to tune the equilibria of membrane proteins within. In this case, we have strong experimental evidence that acyl chain length, and not headgroup chemistry, plays an important role in defining the solvation energetics of the monomeric state of CLC. This is because the solvation structure around the dimerization interface is thinned, twisted and non-bilayer-like introducing an energetic penalty in typical biological membranes containing C16:0,18:1 PO lipid chains. Along these lines, we predict that any lipid of smaller size has the potential to improve the packing and stabilize the free energy of the dissociated state by promoting lipid-lipid interactions and reducing the amount of water penetrating the hydrophobic core. Even using a coarse-grained force-field reflecting a reduced physical model, we find an overall agreement between our calculated values of *ΔΔG* and our experiments. However, the calculations do not recapitulate the energy change that is experimentally observed between 0 → 1% DL, *ΔΔG*_*expt*.,0→1%DL_ = 1.7 ± 0.3 kcal/mole whereas *ΔΔG*_*FEP*, 0→1%DL_ = 0.09 ±0.01 kcal/mole. This could be due to some limitation of the force-field or possibly that our system is of a defined size containing a finite number of DL molecules. Thus, this limits the number of DL that can participate in preferential solvation, whereas in our experiments, the membrane system is practically infinite. Still, the overall agreement of the calculations with experiments has profound implications because it provides a physical model that can be used to now predict the effect of other lipid changes on CLC dimerization equilibria. Note, the effect that we observe does not involve site-specific lipid binding, but rather lipids as non-specific solvent. Yet, this model is completely consistent with our observed experimental change and the linkage relationship, as demonstrated by the similar slope observed in our experiments and FEP calculations. This demonstrates that lipids can finely tune the reactions of membrane proteins via solvation energetics, presenting this as a potential mechanism for how biology is connecting the general membrane protein landscape to physiological changes in lipid composition. Yet, to really understand whether biology readily tunes this driving force will require further experimental and computational investigation in reconstituted and native systems.

This mechanism, where the binding is driven by how the protein changes the shape of the surrounding solvent, is unique to proteins within the membrane. However, it is reminiscent of another important solvent dependent driving force - the hydrophobic effect. In this case, the exposure of non-polar surfaces to polar water restricts the solvent hydrogen bonding network, thereby ordering the water molecules in the first solvation shell. With this, there is a gain in free energy by burying the non-polar surface and then relieving the energetic penalty of the water involved. The broad impact of both driving forces, membrane perturbations and the hydrophobic effect, is that they are non-specific and generalizable. For soluble protein folding and association, the hydrophobic effect provides an innate driving force to bury non-polar groups in a non-specific manner. In the case of membrane protein folding and association, there is an analogous innate driving force to bury surfaces that are disruptive to the membrane bilayer structure. Having a non-specific driving force that favors the condensed state of membrane proteins in membranes is important, as it provides a mechanism for achieving stability while expanding evolutionary sampling of protein sequence. However, a major difference in these two generalizable driving forces is that, in all biological systems, the hydrophobic effect, as the name implies, depends only on water. This is a completely different situation in a biological membrane, which can contain thousands of different types of lipoidal molecules and can readily change in composition due to changes in environmental conditions, metabolic changes, or other physiological situations. Thus, it is conceivable that biology readily tunes this membrane dependent driving force and thus can shift protein equilibria when needed.

Because of the general, and non-specific nature of this driving force, we expect that it will apply not only to oligomerization in membranes, but other equilibrium membrane protein reactions. A recent study on the mechanism of energy-coupling-factor transporters, which involve a substrate binding domain associating with a scaffolding domain, also suggests a mechanism of binding via a thinning of the membrane, promoting the rotation of the substrate domain during association (Faustino et al. 2020). In the case of multi-helix protein folding, helices that expose partially polar regions to the surrounding membrane will introduce local defects that can drive the helices to assemble such that the defect is minimized, where specific protein interactions can then be samples. Along these lines, membrane embedded chaperones and insertases have also shown the ability to introduce thinned defects in the surrounding membrane, suggesting a non-specific mechanism of affinity for un-partnered helices (Wu and Rapoport 2021). In fact, any membrane protein undergoing a conformational change that introduces a perturbation to the surrounding membrane will be linked to the membrane energetics via this mechanism. Certain transport proteins, especially those utilizing the elevator mechanism, introduce large deformations in the membrane with the movement of the transport domain defining the states that provide alternate access for substrates (Zhou et al. 2019; Wang and Boudker 2020). The equilibrium is therefore expected to be linked to the energetic cost of deforming the membrane in both states. In addition, this effect is expected to pertain to the gating of ion channels, and certainly mechanosensitive channels that are already known to be sensitive to the physical state of the membrane and related to the significant distortions of the membrane observed in different gated states. Therefore, the structure of the membrane around various membrane proteins may be an important but often unresolved component when considering the energetic landscape pertaining to folding, binding and conformational behavior.

## Materials and Methods

### Protein purification, fluorophore labeling and reconstitution into lipid bilayers

Methods for protein purification, labeling and reconstitution were carried out as described previously (Chadda et al. 2021; Chadda et al. 2016). CLC-ec1 on the background of C85A/H234C was used in the studies to provide subunit specific labeling. A C-terminal hexahistidine tag is present for purification and is left intact in these studies. Protein was labeled with Cy5-maleimide (sulfo-Cy5, GE biosciences Inc.), with a labeling efficiency of ≈ 0.7 per subunit. CLC-ec1-Cy5 was reconstituted into POPC or 1:4 DLPC/POPC membranes as described previously (Chadda et al. 2016), where the dialysis was carried out at 4 °C for ∼ 3 days, unless otherwise noted.

### Membrane freeze-thaw fusion

Fusion of 2:1 POPE/POPG, POPC and 1:4 DLPC/POPC membranes was monitored by Förster Resonance Energy Transfer in membranes detected by doping vesicles with donor phosphatidyl-ethanolamine-NBD (PE-NBD) and acceptor phosphatidyl-ethanolamine-RhB (PE-RhB) lipids, following the procedures described previously (Cliff et al., 2019). Measurements were conducted using a Fluorolog 3-22 Fluorometer (Horiba, Ltd.).

### Single-molecule photobleaching analysis

After dialysis, liposomes were freeze-thawed five times by cycling through -80 °C and room temperature. The fused membranes were then supplemented with 0.02% NaN_3_ and incubated at room temperature, in the dark, for 4-8 days. The samples were extruded through a 0.4 μm Avestin extruder 21x before examination by single-molecule microscopy. Lanes on the slide were coated with 1 mg/mL poly-lysine and dilute glutaraldehyde to immobilize protein molecules and reduce lateral diffusion. Photobleaching probability distributions were measured as described previously (Chadda et al., eLife 2016; Chadda & Robertson, MIE 2016).

### Equilibrating a coarse-grained POPC bilayer

A pure 1-palmitoyl-2-oleoyl-sn-glycero-3-phosphocholine (POPC) lipid bilayer was built using the insert membrane (insane) python script (Wassenaar et al. 2015) with a simulation box size of 25 nm x 15 nm x 10 nm, including water (90% regular and 10% antifreeze water) and 0.15 M NaCl. Martini 2.0 force field parameters (Marrink et al. 2007) were used to calculate the interactions between coarse-grained beads of the system. The membrane system was energy-minimized and equilibrated with the protocol as described previously (de Jong et al. 2016) and simulated for 20 μs at 303 K and 1 bar with a time-step of 20 fs. The snapshot at the end of 20 μs-long trajectory was extracted to be used to build the CLC-ec1-membrane simulation systems.

### Building a series of coarse-grained CLC-ec1 structural models

The initial atomic coordinates of the wild-type CLC-ec1 dimer (residues 30-458) were extracted from the crystal structure (PDB ID:1OTS) (Dutzler, Campbell, and MacKinnon 2003). The protein structure was converted to a coarse-grained representation with a modified version of the Martinize script (de Jong et al. 2013). The residues E113 in both subunits and D417 in only one subunit were protonated (Faraldo-Gómez and Roux 2004). N and C termini were neutralized. In addition to Martini 2.2(Monticelli et al. 2008) force field parameters used for the protein beads, the overall fold of the protein was maintained with an elastic network having the force constant of 500 kJ/nm^2^/mole and the upper (lower) elastic cut-off distance of 0.9 nm (0.5 nm). This elastic network was built for each protomer separately, without any inter-subunit elastic bonds. The CLC-ec1 dimer built from the crystal structure (PDB ID: 1OTS) was placed so that the dimerization interface is normal to the x axis, while the transmembrane part is parallel to the z axis. This configuration was used for the first umbrella sampling window for the free-energy calculations (window index = 1). Starting from that, we translated the coordinates of one CLC-ec1 protomer 0.05 nm away from the other protomer along the x axis to generate the configuration for the next umbrella sampling window until a separation of 5.5 nm between two protomers was achieved. This resulted in a total of 111 different configurations of two CLC-ec1 protomers, each one being an umbrella window indexed from 1 to 111.

### Building a set of coarse-grained CLC-ec1-membrane systems for the native dimer

Using an in-house MDAnalysis script (Gowers et al. 2019; Michaud-Agrawal et al. 2011), each of the coarse-grained CLC-ec1 protomers built as explained in the previous section was embedded at the center of the preequilibrated POPC bilayer that was extracted from the 20 *μ*s-long coarse grained molecular dynamics simulation as described above. Non-protein beads (lipids, water, and salt ions) within 3 Å of the protein beads were removed to avoid clashes. The net charge of the resulting system was neutralized. Martini 2.0 (Marrink et al. 2007) and Martini 2.2 (Monticelli et al. 2008) force field parameters were used for the non-protein and protein beads, respectively. The coarse-grained protein-membrane system was then equilibrated using the stepwise equilibration protocol suggested in a work conducted by de Jong and coworkers (de Jong et al. 2016). A detailed description of this protocol is presented in **Supp. Table 1**.

The first minimization, with a soft-core potential, was necessary to get rid of the initial steric clashes in the systems and was followed by another minimization with the regular (12-6) Lenard-Jones potential. At the end of these minimization steps, the forces did not get below F_max_ (set to 10 kJ/nm/mole). However, minimization with more steps could cause large gaps between the protein and lipid beads as the lipid molecules attract each other faster than the protein does. Therefore, we continued with the first step of equilibration in which the positions of the backbone beads of the protein were restrained using a harmonic potential with a force constant of 1000 kJ/mole/nm^2^ for 10 ps and then with a force constant of 500 kJ/mole/nm^2^ for 3 ns, while the time step was gradually increased from 2 fs to 20 fs. The pressure and temperature were set to 1 bar and 310 K using the semi-isotropic Berendsen barostat (Berendsen et al. 1984) along with the velocity-rescaling algorithm (Bussi, Donadio, and Parrinello 2007).

A 10 μs-long coarse-grained simulation was carried out for each umbrella window using Gromacs 2019.6 (Abraham et al. 2015). During this production step, we switched to the Parinello-Rahman barostat (Parrinello and Rahman 1981) as suggested by de Jong and coworkers (de Jong et al. 2016) and we used a set of collective variables (1) to keep two CLC-ec1 protomers at the center of the simulation box; (2) to restrain the dimerization interface of one protomer with respect to that of the other protomer; and (3) to restrain the inter-subunit distance between two CLC-ec1 protomers. The harmonic restraints were applied on these collective variables with a force constant of 2000 kJ/mole/nm^2^ using Plumed 2 (Tribello et al. 2014). The detailed description of the collective variables can be found in **Supp. Fig. 3** and **Supp. Table 2**. To keep the protomers at the center of the simulation box, the geometric center of all four G_1_ and G_2_ collective variables was computed for the native and non-native dimers; the *x* and *z* coordinates of this center were restrained using harmonic potentials with a force constant of 2000 kJ/mole/nm^2^. To ensure that the dimerization interface of one protomer faces toward that of the other protomer, the *y* coordinates of the geometric centers of G_1_ and G_2_ collective variables of each protomer were restrained to their initial values based on the 1OTS crystal structure using harmonic potentials with a force constant of 2000 kJ/mole/nm^2^. This way, the protomers always faced each other as intended throughout the simulations, and they were free to reorient themselves within the lipid bilayer. Finally, the inter-subunit distance was defined using a distance vector pointing from the geometric center of G_1_ and G_2_ collective variables of one protomer to that of the other protomer. The *x* coordinate of this distance vector was restrained to fixed values from the native (or non-native) dimer configuration to two dissociated protomers separated by 5.5 nm through umbrella windows indexed 1 to 111, with an increment of 0.05 nm. For this, the range of inter-subunit distance values covered was [3.4 nm, 8.9 nm]. A harmonic bias potential with a force constant of 2000 kJ/mole/nm^2^ to keep the inter-subunit distance at a given value.

### Building a set of coarse-grained CLC-ec1-membrane systems for the non-native dimer

We used the HDOCK server (Yan et al. 2020) to dock two CLC-ec1 protomers in order to obtain a non-native dimer model in which the native dimerization interfaces are exposed to the membrane. Within the top 20 results, we chose a non-native dimer model in which the same non-native interface of two CLC-ec1 protomers face toward each other and their orientation with respect to the membrane is close to that of the native dimer. We then converted the non-native dimer to a coarse-grained representation following the same protocol that we applied to the native dimer. The elastic network was added to keep the overall fold of each monomer, without any inter-subunit elastic bonds. The non-native dimer was also separated starting from its dimer configuration to a separation of 5.5 nm along the *x* axis, creating a total of 111 configurations of two CLC-ec1 protomers through which the inter-subunit distance changed from 3.8 nm to 9.3 nm. These systems were then embedded into the same pre-equilibrated POPC bilayer, and the protein-membrane systems were equilibrated using the stepwise equilibration protocol as described earlier. At the end of the last equilibration step, we defined the same two collective variables (G_1_ and G_2_) per each protomer using a different subset of amino acids as presented in **Supp. Table 2** and carried out a 10 μs-long production run with the same simulation parameters we used for the native dimer for each of 111 umbrella windows.

### Building a set of coarse-grained CLC-ec1-membrane systems to investigate the effect of shorter-chain lipids

To inspect the effect of shorter-chain lipids, we repeated the simulations of two CLC-ec1 with a lipid bilayer composed of 70% 1-palmitoyl-2-oleoyl-sn-glycero-3-phosphocholine (POPC) and 30% 1,2-dilauroyl-sn-glycero-3-phosphocholine (DLPC) lipids. First, we extracted last frames of 10 μs-long production trajectories of the native dimer systems and removed the periodic boundary effects using the trjconv tool of Gromacs (Abraham et al. 2015). Then 30% of the lipids from each leaflet were randomly selected and converted to DLPC. Using the same set of collective variables, we carried out 40 μs-long production runs starting with random velocities at 310 K and 1 bar for a set of 111 umbrella windows while keeping the rest of the simulation parameters identical to the native and non-native dimer models. The covered range for the inter-subunit distance is identical to the native dimer model embedded in a pure POPC bilayer: [3.4 nm, 8.9 nm].

### Computing the free energy contribution of the membrane in CLC-ec1 dimerization

We combined the inter-subunit distance data at every 500 ps extracted from the production runs of all 111 umbrella windows. We calculated the total number of inter-subunit contacts with a distance cutoff of 0.6 nm with the coordination number collective variable for all umbrella windows. In this computation, we set two groups of collective variables composed of the first side chain bead (SC1) of a subset of amino acids located at the membrane-facing transmembrane helices of each CLC-ec1 protomer. The subset of amino acids is listed in **Supp. Table 3**. All other parameters for the coordination number collective variable were set to their default values. We then used a sigmoidal equation to relate the inter-subunit distance to the number of inter-subunit contacts with the following equation:

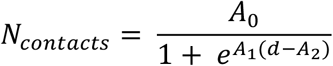

Here, *N*_*contacts*_ and *d* represent the number of inter-subunit contacts and the inter-subunit distance, respectively. *A*_0_, *A*_1_, and *A*_2_ were set to 200, 0.31 and 32.3 for the native dimer (100 POPC), 200, 0.35 and 36.4 for the non-native dimer (100% POPC) and 200, 0.30 and 33.1 for the native dimer (30% DLPC, 70% POPC). The resulting correlation coefficients were 0.99, 0.98 and 0.99 for the native (100 POPC), non-native (100 POPC) and native dimer (30% DLPC, 70% POPC), respectively. This way, we determined the inter-subunit distance cut-off value at which 85% of maximum inter-subunit contacts were lost for each system: 4.05 nm for the native dimer in a POPC membrane, 4.37 nm for the non-native dimer in a POPC membrane and 4.15 nm for the native dimer in a 30% DLPC, 70% POPC membrane.

Next, we used the force-correction analysis method (FCAM) (Marinelli and Faraldo-Gómez 2021) and computed the free energy as a function of inter-subunit distance. FCAM bins each snapshot of the trajectories obtained from umbrella sampling simulations and computes the Boltzmann weight of each bin. To compute the membrane contribution to the free energy in CLC-ec1 dimerization, we reweighted the Boltzmann weight of each frame by multiplying the original Boltzmann weight of the frame with a repulsive harmonic potential having a force constant of 15.4 kcal/mole/Å^2^. The repulsive harmonic potential is set to zero when the inter-subunit distance becomes equal and smaller than the cut-off value at which 85% of maximum inter-subunit contacts were lost for each system. In other words, we computed the free energy profile as a function of inter-subunit contacts as if the simulations were run with a repulsive potential that does not allow two protomers establish more than 15% of potential maximum contacts. Since the protomers are not allowed to maintain a significant amount of inter-subunit contacts, it is the membrane’s contribution that effectively dominates the resulting free energy profile as a function of inter-subunit distance between two protomers.

### Free Energy Perturbation simulations

FEP simulations were based on the CLC-ec1 crystal structure deposited to the protein data bank as entry 1OTS (Dutzler, Campbell, and MacKinnon 2003). This structure was used for simulations of the dimer and the protomers were translated to achieve a separating distance of 5.5 nm for the simulations of the monomers. Following our previous studies, each protomer was truncated up to residue 30. The protein was then converted into a coarse-grained model using the Martinize tool and the membrane and solvent were built around the protein using the insane script. The net charge was neutralized, and the system buffered to 0.150 M with Na^+^ and Cl^-^ ions. An energy minimization step was then performed, and the system equilibrated at constant pressure and temperature for 10 ns. This resulted in a box with approximate dimensions 25.2 nm x 15.1 nm x 9.6 nm. The coordinates of this system were used to create the initial structure for each transformation by modifying select residue types in the topology file; see **Table 4** for details. Each system was then equilibrated for 1 *μ*s at constant temperature and pressured. This equilibration step was followed by a slow growth simulation where the transformation was performed over the course of 20 *μ*s. To facilitate the alchemical transformation, a DLPC* molecule was created that is identical to DLPC but with an extra bead attached to each alkyl chain (**Fig. 4C**). These added beads were given standard bonding and angle bending parameters (**Supp. Table 6**), and their Lennard-Jones terms were set to nil. The free energy perturbation simulations were then carried out using Hamiltonian replica exchange such that lambda was varied for each replica. Lambda values were selected so that the relative entropy between neighboring replicas was maintained at 1 kT or less, see **Supp. Fig. 6**. This requirement resulted in 38 replicas for the 0-1% transformation and 93 for the remaining; the lambda values for each are listed in **Table 5**. The initial conditions for each replica were extracted from the slow growth trajectory such that starting configurations were in equilibrium with the replica’s lambda value. Production runs were carried out with GROMACS 2018.8 using an integration step of 20 fs and exchange moves between neighboring replicas were attempted every 1 ns. This resulted in an exchange rate of approximately 40% on average. The pressure and temperature were set to 1 bar and 303 K using the semi-isotropic Berendsen barostat (Berendsen et al. 1984) along with the velocity-rescaling algorithm(Bussi, Donadio, and Parrinello 2007).The electrostatics were handled using the reaction-field method. To enable error estimates, independent FEP simulations were performed for each transformation such that the forward and reverse pathways were explored. Trajectories were collected using 6 *μ*s for each replica for the 1→10%, 10→20% and 20→30% DLPC conditions and 40 *μ*s for the 0→1% condition. A total of 12.7 ms of simulation time was acquired.

### Error Estimations for Free Energy Perturbation simulations

*ΔG* values are reported for each transformation as the average over the forward and backward paths such that:

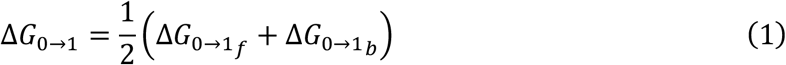

and the uncertainty is given by the standard deviation:

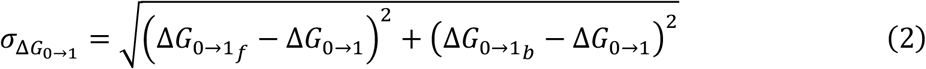

The free energy change over the full transformation is then given as the sum of the individual steps:

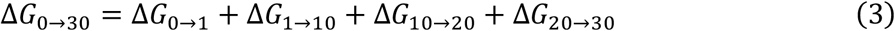

and the uncertainty is computed using standard error propagation methods such that:

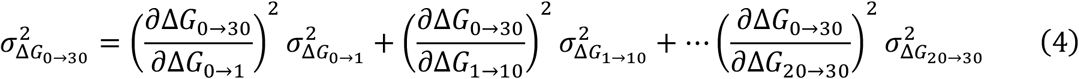

Since the partial derivatives in equation 4 are each unity, this simplifies to:

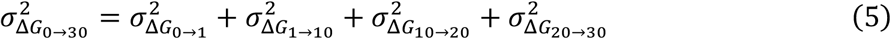

It then follows that ΔΔ,_0→40_is computed as:

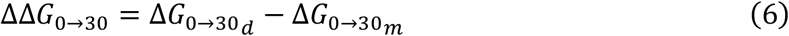

where we have added the subscripts d and m to indicate that the transformation was performed for the dimer or a system containing a pair of monomers. The uncertainty is computed as:

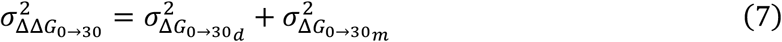

Similarly, the standard error of the mean is given by

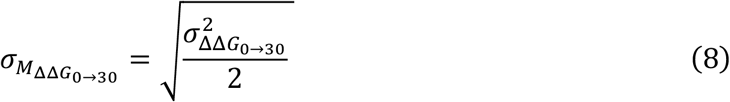

We note that, while the examples shown here focus on the complete transformation, the equations are easily adaptable to partial transformation like 0→1%, 0→10% or 0→20%.

## Notes

### Competing Interest Statement

The authors have declared no competing interest.

## References

Abraham, Mark James, Teemu Murtola, Roland Schulz, Szilárd Páll, Jeremy C Smith, Berk Hess, and Erik Lindahl. 2015. ‘GROMACS: High performance molecular simulations through multi-level parallelism from laptops to supercomputers’, SoftwareX, 1: 19–25.

Aleksandrova, Antoniya A., Edoardo Sarti, and Lucy R. Forrest. 2020. ‘MemSTATS: A Benchmark Set of Membrane Protein Symmetries and Pseudosymmetries’, Journal of Molecular Biology, 432: 597–604.

Anselmi, Claudio, Karen M Davies, and José D Faraldo-Gómez. 2018. ‘Mitochondrial ATP synthase dimers spontaneously associate due to a long-range membrane-induced force’, Journal of General Physiology, 150: 763–70.

Berendsen, Herman JC, JPM van Postma, Wilfred F Van Gunsteren, ARHJ DiNola, and Jan R Haak. 1984. ‘Molecular dynamics with coupling to an external bath’, The Journal of chemical physics, 81: 3684–90.

Bussi, Giovanni, Davide Donadio, and Michele Parrinello. 2007. ‘Canonical sampling through velocity rescaling’, The Journal of chemical physics, 126: 014101.

Chadda, R., and J. L. Robertson. 2016. ‘Chapter Three - Measuring Membrane Protein Dimerization Equilibrium in Lipid Bilayers by Single-Molecule Fluorescence Microscopy.’ in Maria Spies and Yann R. Chemla (eds.), Methods in Enzymology (Academic Press).

Chadda, Rahul, Nathan Bernhardt, Elizabeth G Kelley, Susana CM Teixeira, Kacie Griffith, Alejandro Gil-Ley, Tuğ ba N Öztürk, Lauren E Hughes, Ana Forsythe, and Venkatramanan Krishnamani. 2021. ‘Membrane transporter dimerization driven by differential lipid solvation energetics of dissociated and associated states’, Elife, 10: e63288.

Chadda, Rahul, Lucy Cliff, Marley Brimberry, and Janice L Robertson. 2018. ‘A model-free method for measuring dimerization free energies of CLC-ec1 in lipid bilayers’, Journal of General Physiology, 150: 355–65.

Chadda, Rahul, Venkatramanan Krishnamani, Kacey Mersch, Jason Wong, Marley Brimberry, Ankita Chadda, Ludmila Kolmakova-Partensky, Larry J. Friedman, Jeff Gelles, and Janice L. Robertson. 2016. ‘The dimerization equilibrium of a ClC Cl-/H+ antiporter in lipid bilayers’, Elife, 5: e17438.

Chadda, Rahul, Taeho Lee, Priyanka Sandal, Robyn Mahoney-Kruszka, and Janice L. xsRobertson. 2023. ‘A thermodynamic analysis of CLC transporter dimerization in lipid bilayers’, bioRxiv: 2023.03.14.532678.

de Jong, Djurre H, Svetlana Baoukina, Helgi I Ingólfsson, and Siewert J Marrink. 2016. ‘Martini straight: Boosting performance using a shorter cutoff and GPUs’, Computer Physics Communications, 199: 1–7.

de Jong, Djurre H., Gurpreet Singh, W. F. Drew Bennett, Clement Arnarez, Tsjerk A. Wassenaar, Lars V. Schäfer, Xavier Periole, D. Peter Tieleman, and Siewert J. Marrink. 2013. ‘Improved Parameters for the Martini Coarse-Grained Protein Force Field’, Journal of chemical theory and computation, 9: 687–97.

Dutzler, Raimund, Ernest B. Campbell, and Roderick MacKinnon. 2003. ‘Gating the Selectivity Filter in ClC Chloride Channels’, Science, 300: 108–12.

Ernst, Melanie, Esam A Orabi, Randy B Stockbridge, José D Faraldo-Gómez, and Janice L Robertson. 2023. ‘Dimerization mechanism of an inverted-topology ion channel in membranes’, bioRxiv: 2023.01.27.525942.

Faraldo-Gómez, José D, and Benoit Roux. 2004. ‘Electrostatics of ion stabilization in a ClC chloride channel homologue from Escherichia coli’, Journal of molecular biology, 339: 981–1000.

Faustino, Ignacio, Haleh Abdizadeh, Paulo C. T. Souza, Aike Jeucken, Weronika K. Stanek, Albert Guskov, Dirk J. Slotboom, and Siewert J. Marrink. 2020. ‘Membrane mediated toppling mechanism of the folate energy coupling factor transporter’, Nature communications, 11: 1763.

Gowers, Richard J, Max Linke, Jonathan Barnoud, Tyler John Edward Reddy, Manuel N Melo, Sean L Seyler, Jan Domanski, David L Dotson, Sébastien Buchoux, and Ian M Kenney. 2019. “MDAnalysis: a Python package for the rapid analysis of molecular dynamics simulations.” In.: Los Alamos National Lab.(LANL), Los Alamos, NM (United States).

Marinelli, Fabrizio, and José D Faraldo-Gómez. 2021. ‘Force-Correction Analysis Method for Derivation of Multidimensional Free-Energy Landscapes from Adaptively Biased Replica Simulations’, Journal of chemical theory and computation, 17: 6775–88.

Marrink, Siewert J., H. Jelger Risselada, Serge Yefimov, D. Peter Tieleman, and Alex H. de Vries. 2007. ‘The MARTINI Force Field: Coarse Grained Model for Biomolecular Simulations’, The Journal of Physical Chemistry B, 111: 7812–24.

Michaud-Agrawal, Naveen, Elizabeth J Denning, Thomas B Woolf, and Oliver Beckstein. 2011. ‘MDAnalysis: a toolkit for the analysis of molecular dynamics simulations’, Journal of computational chemistry, 32: 2319–27.

Monticelli, Luca, Senthil K. Kandasamy, Xavier Periole, Ronald G. Larson, D. Peter Tieleman, and Siewert-Jan Marrink. 2008. ‘The MARTINI Coarse-Grained Force Field: Extension to Proteins’, Journal of chemical theory and computation, 4: 819–34.

Mueller, Benjamin K., Sabareesh Subramaniam, and Alessandro Senes. 2014. ‘A frequent, GxxxG-mediated, transmembrane association motif is optimized for the formation of interhelical Cα–H hydrogen bonds’, Proceedings of the National Academy of Sciences, 111: E888–E95.

Parrinello, Michele, and Aneesur Rahman. 1981. ‘Polymorphic transitions in single crystals: A new molecular dynamics method’, Journal of Applied physics, 52: 7182–90.

Smith, Steven O., Markus Eilers, David Song, Evan Crocker, Weiwen Ying, Michel Groesbeek, Guenter Metz, Martine Ziliox, and Saburo Aimoto. 2002. ‘Implications of Threonine Hydrogen Bonding in the Glycophorin A Transmembrane Helix Dimer’, Biophysical journal, 82: 2476–86.

Tribello, Gareth A, Massimiliano Bonomi, Davide Branduardi, Carlo Camilloni, and Giovanni Bussi. 2014. ‘PLUMED 2: New feathers for an old bird’, Computer Physics Communications, 185: 604–13.

Wang, Xiaoyu, and Olga Boudker. 2020. ‘Large domain movements through the lipid bilayer mediate substrate release and inhibition of glutamate transporters’, Elife, 9: e58417.

Wassenaar, Tsjerk A, Helgi I Ingólfsson, Rainer A Böckmann, D Peter Tieleman, and Siewert J Marrink. 2015. ‘Computational lipidomics with insane: a versatile tool for generating custom membranes for molecular simulations’, Journal of chemical theory and computation, 11: 2144–55.

Wu, Xudong, and Tom A Rapoport. 2021. ‘Translocation of proteins through a distorted lipid bilayer’, Trends in cell biology, 31: 473–84.

Yan, Yumeng, Huanyu Tao, Jiahua He, and Sheng-You Huang. 2020. ‘The HDOCK server for integrated protein– protein docking’, Nature Protocols, 15: 1829–52.

Zhou, Wenchang, Giacomo Fiorin, Claudio Anselmi, Hossein Ali Karimi-Varzaneh, Horacio Poblete, Lucy R Forrest, and José D Faraldo-Gómez. 2019. ‘Large-scale state-dependent membrane remodeling by a transporter protein’, Elife, 8: e50576.

