## Supplementary Figures & Tables for "Mitigation of membrane morphology defects explain stability and orientational specificity of CLC dimers"

### The energetic contribution of the membrane to CLC dimerization

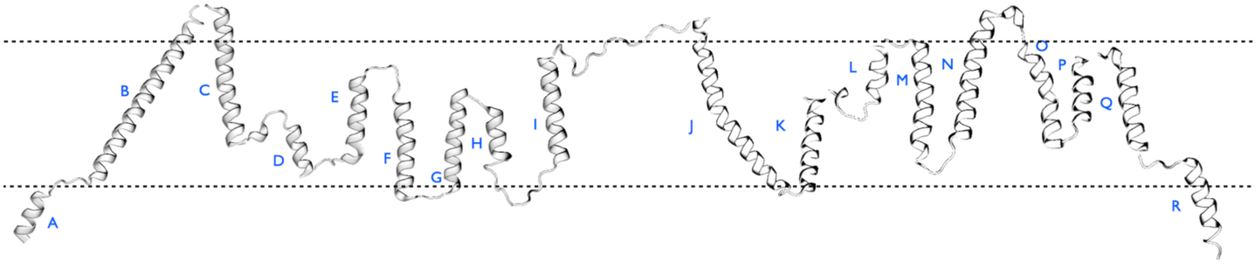


**Supplementary Figure 1.** CLC-ec1 transmembrane helix topology. An un-wound representation of the of CLC-ec1 derived from pdb 1OTS, preserving the transmembrane helical topology. Helices H, I, P and Q form the native dimerization interface.

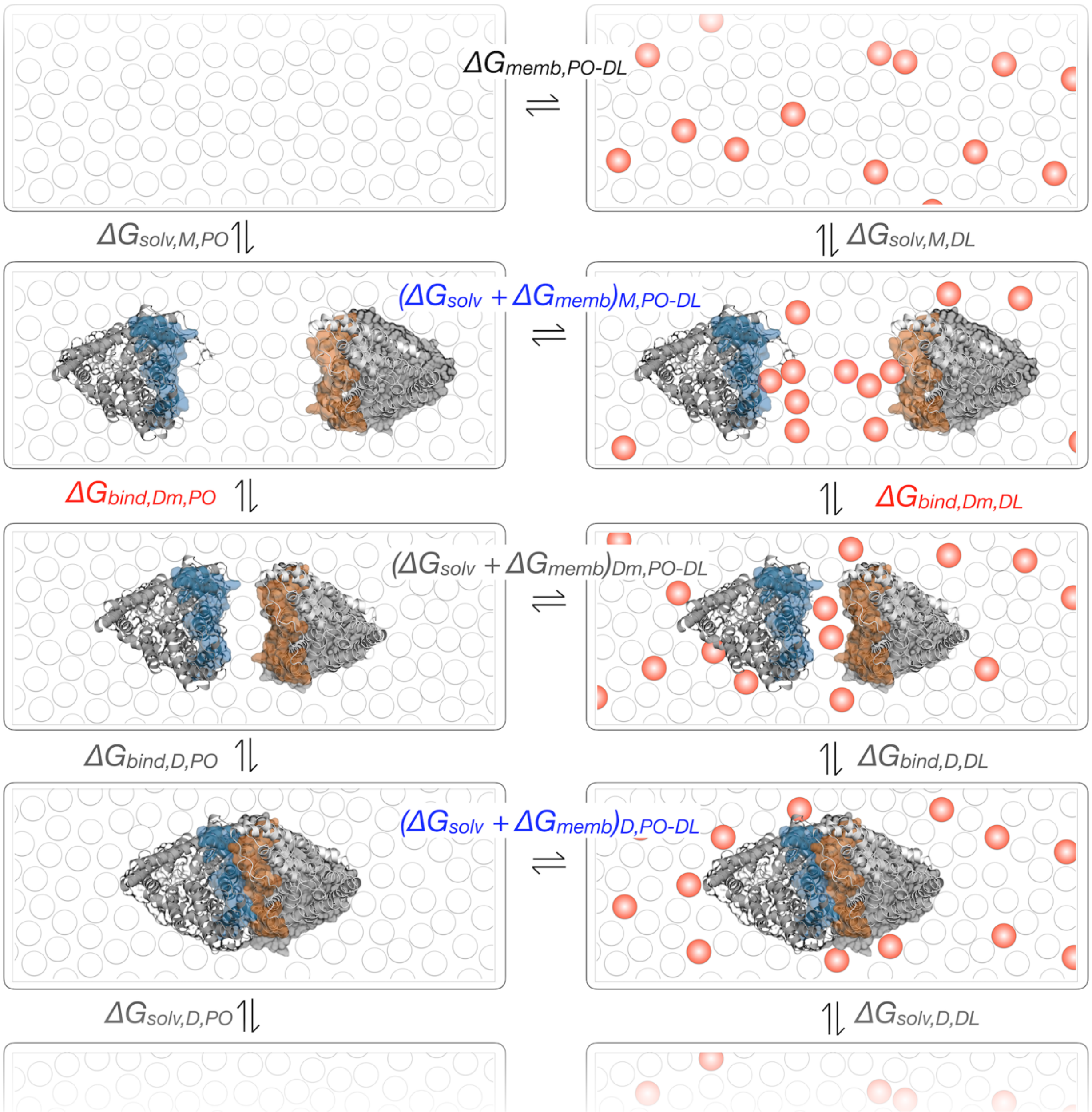


**Supplementary Figure 2. Thermodynamic cycle of CLC dimerization in membranes linked to changes in lipid composition.** *Red*, components of the cycle pertaining to membrane contribution of the potential of mean force associated with binding or dimerization. *Blue*, components of the cycle calculated in the free energy perturbation calculations. *M* – monomeric state, *D* – dimeric state corresponding to the crystal structure, $D_m$ – state where two subunits are closely associated but without protein contacts. The left leg of the reaction indicates the thermodynamic cycle for dimerization in PO lipids (white circles), while the right leg indicates a different lipid composition, here DL (red circles) in PO.

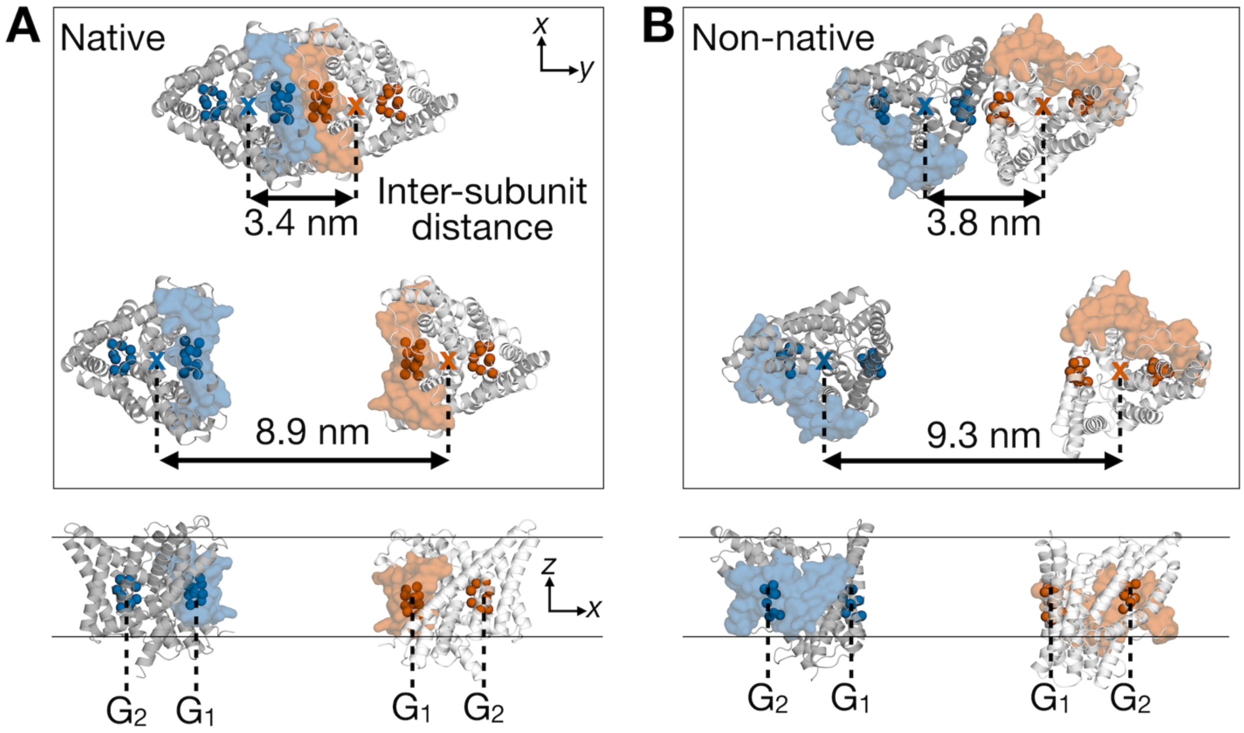


**Supplementary Figure 3. Definition of inter-subunit distance for native and non-native CLC dimers.** (**A**) native and (**B**) non-native dimer models from the extracellular side of the membrane (top) and from within the membrane core (bottom). CLC-ec1 protomers are displayed in cartoon representation, colored white and gray. Four transmembrane helices forming the native dimerization interface are overlaid as a transparent surface in orange and blue. In each protomer, alpha carbons of residues used in the inter-subunit distance calculation are shown, represented as orange or blue spheres. Backbone beads of residues were grouped into four centers ($G_1$ and $G_2$ per promoter) and used for defining the restraints that kept the inter-subunit distance at a fixed value while ensuring that dimerization interfaces of two subunits always face each other. List of these residues for the native and non-native dimer models are presented in **Supp. Table 2.**

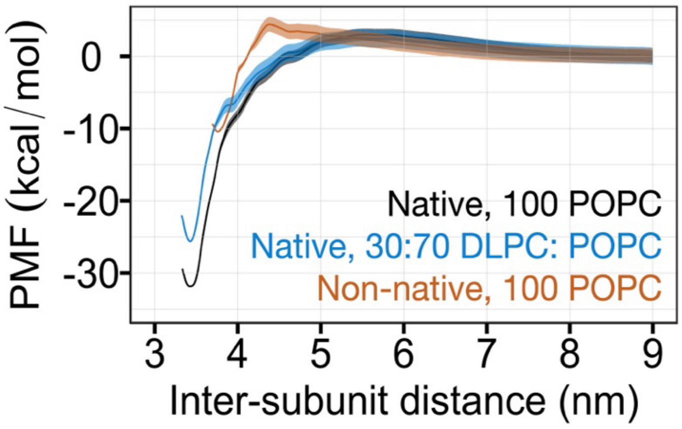


**Supplementary Figure 4. Potential of mean force (PMF) profiles for dimerization of CLC-ec1 protomers as a function of inter-subunit distance.** The PMF profiles were generated using the FCAM and colored as black for the native dimerization interface in a pure POPC bilayer, blue for the native dimerization interface in a 30% DLPC, 70% POPC bilayer and orange for the non-native dimerization interface in a pure POPC bilayer. A series of 111 independent 10 $\mu$s-long coarse-grained molecular dynamics simulations were performed for the systems within pure POPC bilayers and 40 $\mu$s-long simulations for that within a mixture of DLPC and POPC lipids. For all three systems under investigation, the standard error values around the mean PMF values were estimated with block averaging of PMF profiles obtained from four separate parts of these trajectories and plotted as shaded areas.

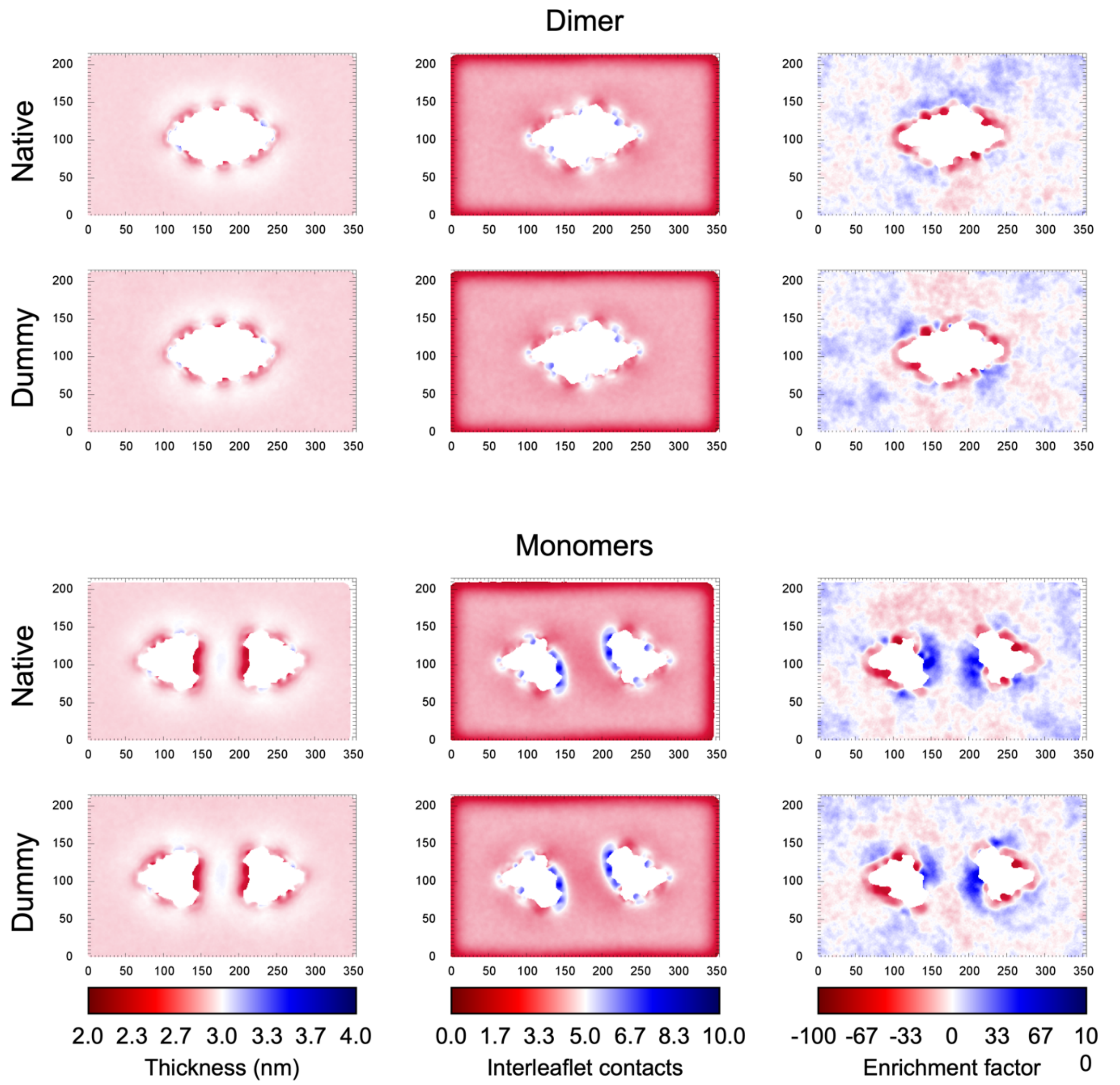


**Supplementary Figure 5.** Comparison of 3 metrics between systems containing a mixture of POPC and native DLPC molecules and systems containing POPC and DLPC*. The left panel compares the membrane thickness for both molecule types. The middle panel compares the number of interleaflet contacts, and the right panel focuses on the DLPC enrichment factor. Results are given for both the dimer (upper panels) and a pair of monomers (lower panels).

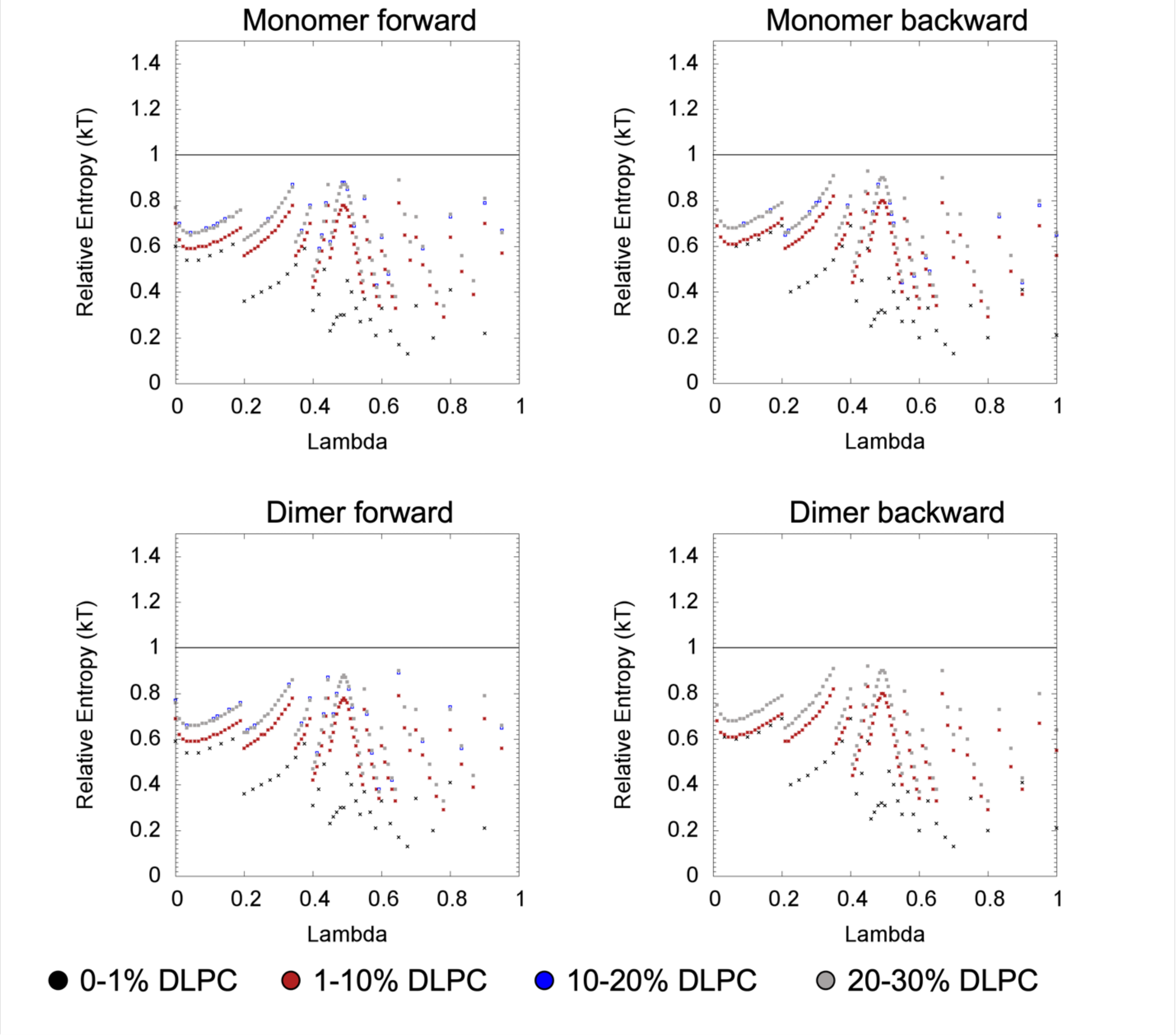


**Supplementary Figure 6.** Relative entropy between the replica, whose lambda value is given in x-axis, and the neighboring replica to the right for the forward path or left for the backward path.

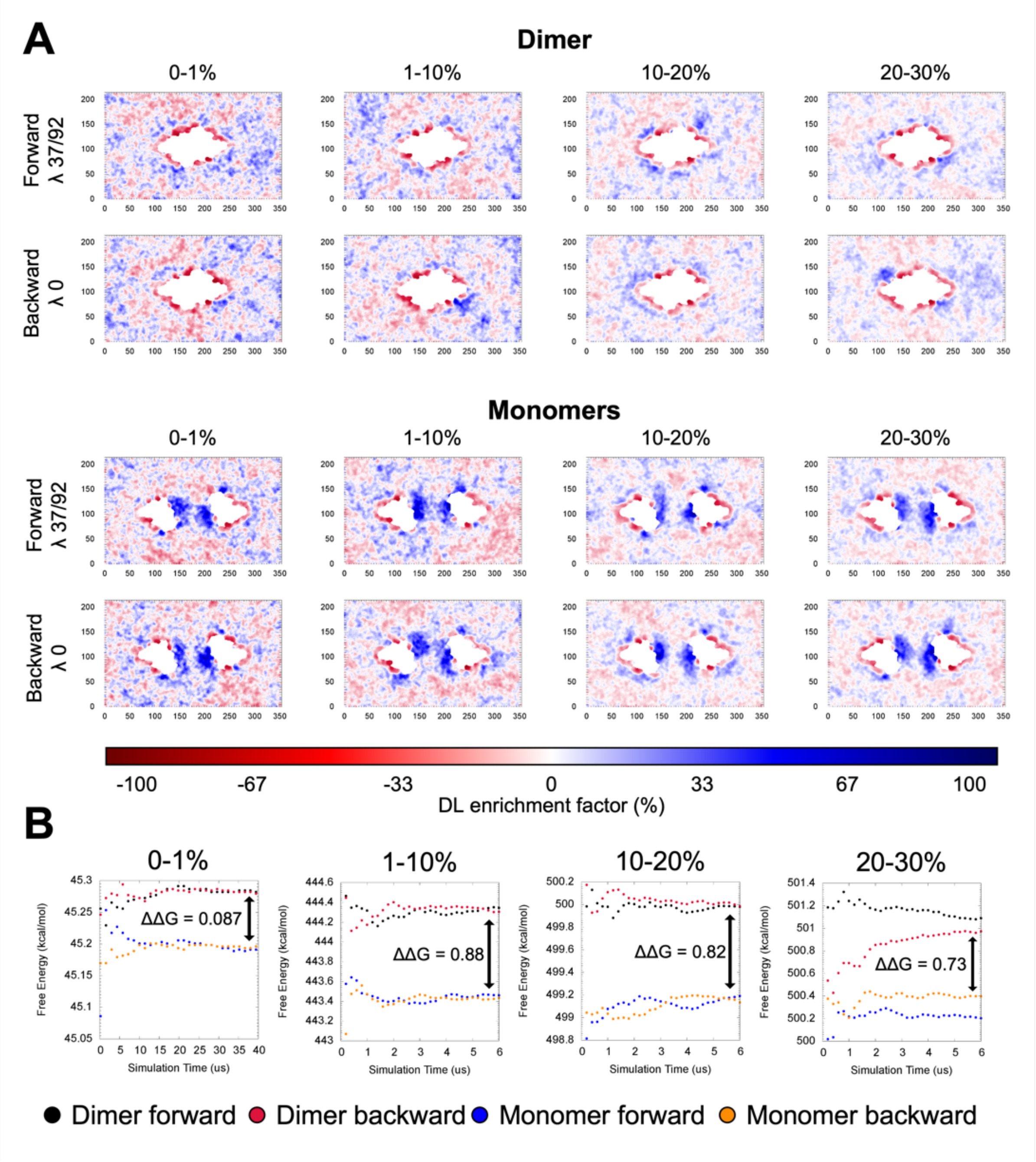


**Supplementary Figure 7. (A)** The DLPC enrichment factor given for select replicas that span each transformation as well as the forward and backward paths. Only replicas whose dynamics are physical are chosen; in particular, we choose the replicas with a lambda value of 1. The simulation conditions are indicated above each plot. Since the lambda value shown contains a lambda value of 1, the plots correspond to 1%, 10%, 20% and 30% DLPC moving left to right. **(B)** The free energy estimate reported for each transformation as a function of the simulation time. *ΔΔG* is also shown for each transformation and is indicated with the arrows. The indicated distance dictates the acceptable error for each transformation. That is, the error must be small for each transformation relative to *ΔΔG* or the signal will be lost.

**Supplementary Table 1.** The stepwise equilibration protocol for the coarse-grained CLC-membrane systems.

| Step | Number of steps | Time step | Force constant (positional restraints on the protein beads) |
|---|---|---|---|
| Minimization 1* | 1000 (steepest descent) | — | — |
| Minimization 2 | 1000 (steepest descent) | — | — |
| Equilibration 1 | 5000 (10 ps) | 2 fs | 1000 kJ/$nm^2$/mol |
| Equilibration 2 | 500000 (1 ns) | 2 fs | 500 kJ/$nm^2$/mol |
| Equilibration 3 | 100000 (1 ns) | 10 fs | 500 kJ/$nm^2$/mol |
| Equilibration 4 | 50000 (1 ns) | 20 fs | 500 kJ/$nm^2$/mol |

**A soft-core potential was used instead of the 12-6 Lennard-Jones potential.*

**Supplementary Table 2.** The list of amino acids used to define four collective variables for the native and non-native dimers.

| **Native dimer** | | | | **Non-native dimer** | | | |
|---|---|---|---|---|---|---|---|
| **Protomer 1** | | **Protomer 2** | | **Protomer 1** | | **Protomer 2** | |
| **$G_1$** | **$G_2$** | **$G_1$** | **$G_2$** | **$G_1$** | **$G_2$** | **$G_1$** | **$G_2$** |
| L194 | T138 | L194 | T138 | F53 | G364 | P129 | G395 |
| A195 | M143 | A195 | M143 | D54 | G393 | V130 | A396 |
| G196 | V144 | G196 | V144 | K55 | M394 | K131 | L397 |
| L198 | L145 | L198 | L145 | G56 | G395 | F132 | A399 |
| F199 | C347 | F199 | C347 | V57 | A396 | F133 | I402 |
| L406 | F348 | L406 | F348 | F132 | L397 | G134 | R403 |
| T407 | A352 | T407 | A352 | F133 | L398 | G135 | A404 |
| G408 | P353 | G408 | P353 | G135 | A400 | L136 | G408 |
| I410 | G354 | I410 | G354 | L136 | | G137 | |
| L411 | | L411 | | | | | |

**Supplementary Table 3.** List of amino acids chosen for each protomer to calculate the number of inter-subunit contacts

| | | | | | | | |
|---|---|---|---|---|---|---|---|
| F37 | L84 | I154 | K216 | L256 | T289 | V338 | Q420 |
| V41 | V88 | I158 | F219 | I260 | L293 | I342 | L423 |
| T44 | F92 | N191 | I223 | I264 | I298 | L346 | L430 |
| L48 | L96 | L194 | I227 | I268 | L301 | P405 | L434 |
| K55 | P129 | I197 | I231 | L274 | L305 | I410 | |
| W59 | F133 | I201 | H234 | L279 | V334 | L413 | |

**Supplementary Table 4.** Number of molecules simulated for each FEP transformation.

| **Molecule** | **0-1 % DLPC** | **1-10 % DLPC** | **10-20 % DLPC** | **20-30 % DLPC** |
|---|---|---|---|---|
| Protein | 2 | 2 | 2 | 2 |
| POPC | 1,082 | 984 | 874 | 764 |
| POPC_to_DLPC | 10 | 98 | 110 | 110 |
| D_DLPC | 0 | 10 | 108 | 218 |
| W | 14,552 | 14,552 | 14,552 | 14,552 |
| WF | 1,616 | 1,616 | 1,616 | 1,616 |
| $Na^+$ | 172 | 172 | 172 | 172 |
| $Cl^-$ | 185 | 185 | 185 | 185 |

**Supplementary Table 5.** λ values used in FEP simulations.

| Monomers/Dimer (0-1 % DLPC) | | | | | | | | | | |
|---|---|---|---|---|---|---|---|---|---|---|
| | **1's** | **10's** | **20's** | **30's** | **40's** | **50's** | **60's** | **70's** | **80's** | **90's** |
| **0** | 0 | 0.3 | 0.48 | 0.625 | | | | | | |
| **1** | 0.03333 | 0.325 | 0.49 | 0.65 | | | | | | |
| **2** | 0.06667 | 0.35 | 0.5 | 0.675 | | | | | | |
| **3** | 0.1 | 0.375 | 0.5125 | 0.7 | | | | | | |
| **4** | 0.13333 | 0.4 | 0.525 | 0.75 | | | | | | |
| **5** | 0.16667 | 0.41667 | 0.5375 | 0.8 | | | | | | |
| **6** | 0.2 | 0.43333 | 0.55 | 0.9 | | | | | | |
| **7** | 0.225 | 0.45 | 0.56667 | 1 | | | | | | |
| **8** | 0.25 | 0.46 | 0.58333 | | | | | | | |
| **9** | 0.275 | 0.47 | 0.6 | | | | | | | |

| Monomers/Dimer (1-10 % DLPC, 10-20 % DLPC, 20-30 % DLPC) | | | | | | | | | | |
|---|---|---|---|---|---|---|---|---|---|---|
| | **1's** | **10's** | **20's** | **30's** | **40's** | **50's** | **60's** | **70's** | **80's** | **90's** |
| **0** | 0 | 0.11111 | 0.22 | 0.32 | 0.40625 | 0.465 | 0.515 | 0.57143 | 0.66667 | 0.9 |
| **1** | 0.01111 | 0.12222 | 0.23 | 0.33 | 0.4125 | 0.47 | 0.52 | 0.57857 | 0.68333 | 0.95 |
| **2** | 0.02222 | 0.13333 | 0.24 | 0.34 | 0.41875 | 0.475 | 0.525 | 0.58571 | 0.7 | 1 |
| **3** | 0.03333 | 0.14444 | 0.25 | 0.35 | 0.425 | 0.48 | 0.53 | 0.59286 | 0.72 | |
| **4** | 0.04444 | 0.15556 | 0.26 | 0.35833 | 0.43125 | 0.485 | 0.535 | 0.6 | 0.74 | |
| **5** | 0.05556 | 0.16667 | 0.27 | 0.36667 | 0.4375 | 0.49 | 0.54 | 0.61 | 0.76 | |
| **6** | 0.06667 | 0.17778 | 0.28 | 0.375 | 0.44375 | 0.495 | 0.545 | 0.62 | 0.78 | |
| **7** | 0.07778 | 0.18889 | 0.29 | 0.38333 | 0.45 | 0.5 | 0.55 | 0.63 | 0.8 | |
| **8** | 0.08889 | 0.2 | 0.3 | 0.39167 | 0.455 | 0.505 | 0.55714 | 0.64 | 0.83333 | |
| **9** | 0.1 | 0.21 | 0.31 | 0.4 | 0.46 | 0.51 | 0.56429 | 0.65 | 0.86667 | |

**Supplementary Table 6.** *Force field parameters for DLPC**

| **Bonding Terms** | | |
|---|---|---|
| ATOMS | LENGTH (NM) | FORCE CONSTANT (KJ MOL$^{-1}$) |
| 1-2 | 0.47 | 1250 |
| 2-3 | 0.47 | 1250 |
| 3-4 | 0.37 | 1250 |
| 3-5 | 0.47 | 1250 |
| 5-6 | 0.47 | 1250 |
| 6-7 | 0.47 | 1250 |
| 7-8 | 0.47 | 1250 |
| 4-9 | 0.47 | 1250 |
| 9-10 | 0.47 | 1250 |
| 10-11 | 0.47 | 1250 |
| 11-12 | 0.47 | 1250 |

| **Angle Terms** | | |
|---|---|---|
| ATOMS | ANGLE (°) | FORCE CONSTANT (KJ MOL$^{-1}$ RAD$^{-2}$) |
| 2-3-4 | 120 | 25 |
| 2-3-5 | 180 | 25 |
| 3-5-6 | 180 | 25 |
| 5-6-7 | 180 | 25 |
| 6-7-8 | 180 | 25 |
| 4-9-10 | 180 | 25 |
| 9-10-11 | 180 | 25 |
